# Non-mydriatic chorioretinal imaging in a transmission geometry and application to retinal oximetry

**DOI:** 10.1101/314765

**Authors:** Timothy D. Weber, Jerome Mertz

## Abstract

The human retina is typically imaged in a reflection geometry, where light is delivered through the pupil and images are formed from the light reflected back from the retina. In this configuration, artifacts caused by retinal surface reflex are often encountered, which complicate quantitative interpretation of the reflection images. We present an alternative illumination method, which avoids these artifacts. The method uses deeply penetrating near-infrared (NIR) light delivered transcranially from the side of the head, and exploits multiple scattering to redirect a portion of the light towards the posterior eye. This unique transmission geometry simplifies absorption measurements and enables flash-free, non-mydriatic imaging as deep as the choroid. Images taken with this new transillumination approach are applied to retinal oximetry.

**OCIS codes:** (170.4460) Ophthalmic optics and devices; (170.2945) Illumination design; (170.1470) Blood or tissue constituent monitoring.

## 1. Introduction

A wealth of information is contained in the optical appearance of the posterior human eye (i.e. ocular fundus) [1]. The retina possesses a tightly regulated blood supply similar to that of the brain [2]. Misregulation or damage to retinal circulation has been implicated in ocular diseases such as age-related macular degeneration (AMD) and systemic diseases such diabetes and cardiovascular disease [3]. Recent work has suggested that possible biomarkers for neurodegenerative conditions, like Alzheimer’s disease, may also exist in the retinal vasculature [4, 5]. Thus, there is strong need for high-throughput, noninvasive methods to monitor retinal circulation.

Retinal imaging methods can be divided into widefield and scanning-based techniques. Scanning techniques raster a focused laser beam across the fundus while recording back-reflected light, and images are synthesized point-by-point. The use of confocal detection [6] or coherence gating [7] enables 3D sectioning and very high contrast by rejecting multiple scattered light that otherwise contributes to background. Indeed, variations on scanning techniques, such as optical coherence tomography angiography [8] and adaptive optics scanning ophthalmoscopy [9], have produced detailed views of retinal capillary networks in vivo.

In contrast, widefield techniques illuminate large areas of the fundus and record real images of the retina from light reflected back through the pupil. In the clinic, this is routinely performed with a fundus camera and generally provides adequate contrast of major structures, such as the vasculature, optic disc, nerve fiber bundle, and macula, depending on the wavelength choice [10]. A significant disadvantage of widefield techniques is that they do not provide depth discrimination. Nevertheless, owing to their simplicity and ubiquity in clinical practice, widefield techniques are more suitable for high-throughput disease screening.

Although exogenous vessel contrast agents, such as fluorescein [11], have been used for decades, in principle, natural spectral differences between endogenous absorbers should reveal the relative distributions of chromophores in the eye [12]. Specifically, quantification of oxygen saturation in retinal vessels is known as retinal oximetry (see Ref. [13] for an introduction, or Ref. [14] for a recent comprehensive review). It has repeatedly been shown that in major ocular diseases, retinal vessel oxygen saturation abnormalities precede clinically detectable morphological changes [15].

The challenge with all spectroscopic techniques in the eye is that multiple reflection paths from stratified fundus layers contribute to the observed reflectance image and that the relative contribution from each of these reflections is highly sensitive to the degree of fundus pigmentation [16]. Models for the interpretation of the reflected light have characterized the return of light according to wavelength [16–18] and refractive index differences between retinal layers [19], but research is still needed to fully characterize the light remitted from ocular fundus structures. Alternatively, oximetry can be performed without a full optical model. Scanning techniques with depth discrimination, such as visible light-OCT [20], are able to isolate just the light paths traversing retinal vessels and provide oximetric data. Widefield approaches based on modified fundus cameras [21–24] also provide oximetric data with the aid of a calibration. However, widefield retinal oximetry is crucially dependent on calibration and is susceptible to artifacts related to specular back reflections (reflex). These artifacts obscure the spectral signatures used to detect chromophores. A well-known artifact is central vessel reflex which appears as bright glints near the center of otherwise absorbing vessels [10, 25]. The reflex is more intense for large arteries, and may appear and disappear repeatedly along the length of the vessel, leading to the so-called “rattlesnake” artifact in oximetric maps [14]. The inner limiting membrane and superficial nerve fiber layer (NFL) of the retina also cause a smoothly varying glare across the fundus, particularly for young eyes. Researchers have gone to extreme lengths to avoid these confounding reflections, even so far as puncturing the eye (of swine) to illuminate the retina from highly oblique angles [26].

In theory, unwanted reflections could be avoided by adopting a transmission imaging geometry. The key difference is that in a reflection geometry, anterior reflection artifacts are first order in strength, whereas in a transmission geometry they are third order: at least two extraneous reflections are required to cause an artifact in the transmission direction. However, it is not immediately obvious how to achieve such a geometry without resorting to unacceptably invasive methods. The solution proposed here is inspired by compelling applications that exploit the capacity of near-infrared (NIR) light to readily penetrate the head and probe oxygen saturation dynamics deep inside the brain [27]. Certainly if enough light reaches the brain (and back) to enable these applications, then it should also be possible to illuminate the back of the eye with NIR light.

Specifically, we propose an alternative widefield fundus imaging strategy based on light delivered transcranially through the subject’s temple. The light diffuses through the bone and illuminates the retina not from the front, as in standard techniques, but rather mostly from the back. As such, images are formed from light *transmitted* through the retina rather than *reflected* from the retina. In this way, the image formation is conceptually similar to that of oblique back-illumination microscopy [28], which is capable of transmission-like phase-gradient and absorption contrast microscopic imaging in thick tissue. The use of NIR light also permits imaging without mydriatics or uncomfortable flash exposures.

The purpose of this paper is to communicate our preliminary results obtained using this new transmission fundus imaging method. The system design and image processing methods are described in detail. Finally, we present an example of retinal oximetry using the technique.

## 2. Methods

### 2.1 Fundus transillumination imaging system

The system consists of illumination and detection subsystems shown together in Fig. 1A and detailed separately below. An example raw transcranial fundus image is provided in Fig. 1B.

**Fig. 1.**
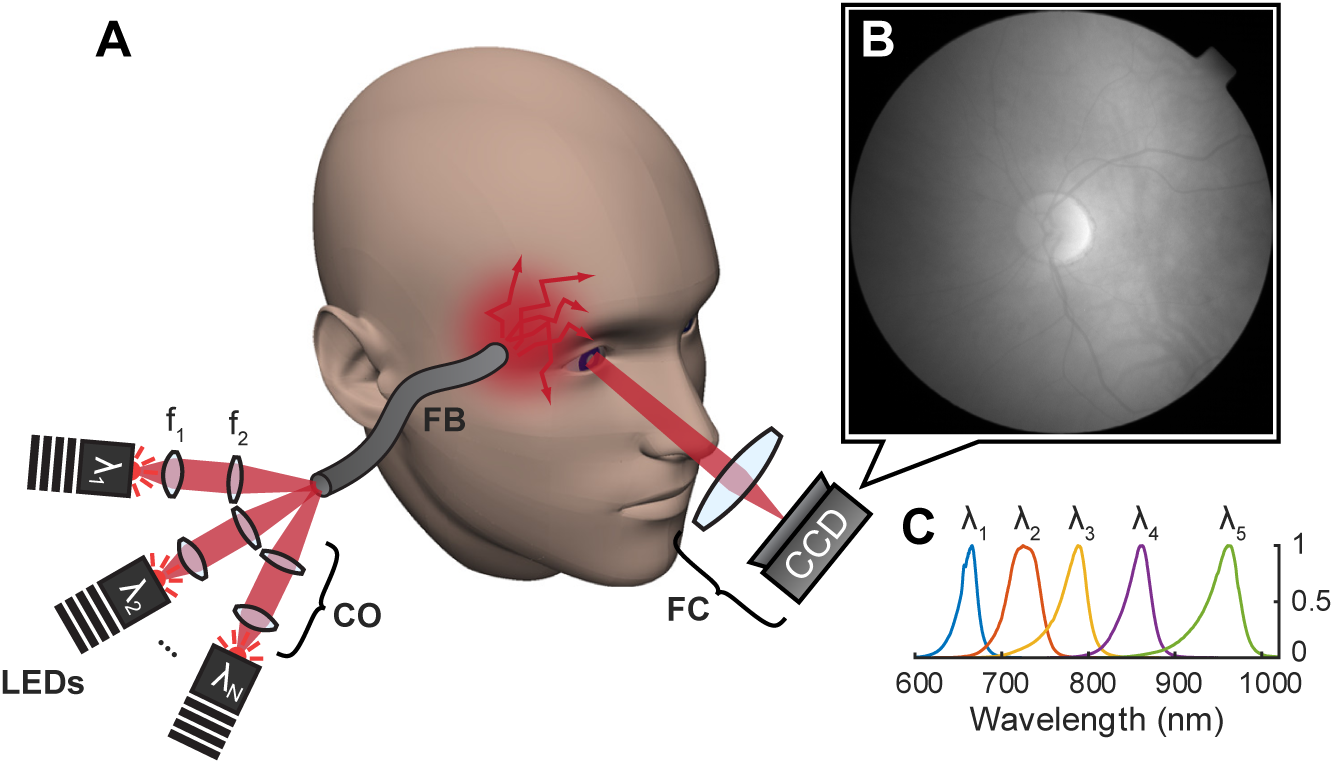
A: Schematic of fundus transillumination and imaging. LEDs at several central wavelengths (λ_N_) are imaged via coupling optics (CO), comprised of lenses f_1_ and f_2_, onto the proximal end of a flexible fiber bundle (FB). A commerical fundus camera (FC) images the transilluminated fundus onto a camera (CCD). B: Example raw image recorded on the CCD. C: Normalized measured spectra of available high-power deep red and NIR LEDs.

#### 2.1.1 Transcranial illumination

According to the theory of diffuse light transport [29], only a small fraction of the total light available is expected to penetrate centimeters through the head and illuminate the posterior eye. LEDs represent a cost-effective and scalable means of obtaining the necessary light power at several wavelengths in the NIR spectrum. Five high-power LEDs ranging in center wavelength from 660 to 940 nm were used to provide transcranial illumination (detailed in Table 1). When active, an LED driver (Thorlabs, LEDD1B) applied the maximum specified current to the particular LED. Resulting spectra are shown in Fig. 1C. To avoid possible subject discomfort resulting from direct contact with hot LED PCBs, the light was delivered remotely through a 13 mm-diameter flexible fiber bundle. Each LED was imaged onto the proximal end of the fiber bundle (f_1_ = 16 mm, Thorlabs ACL25416U-B; f_2_ = 60 mm, Thorlabs LA11340-B). The distal end of the bundle was gently pressed onto the skin near the subject’s temple. With careful (3-dimensional) alignment, up to 6 LEDs could be coupled into the bundle, with each LED beam path sharing a portion of the bundle’s distal end acceptance cone. In this way, different combinations of LEDs (or several at the same wavelength) could be simultaneously active, however this feature was not used in the present study.

After convergence and transmission through the fiber bundle, a power meter was used to measure irradiance to compare with the ANSI Z136.1 safety standard [30]. The measured irradiance at each wavelength was equal to or below the standard’s maximum permissible exposure (MPE) for skin (see Table 1). Continuous exposure (i.e. >10 sec) was assumed. The spectral full width at half maximum (FWHM) for each LED was small enough that only the LED center wavelength was considered. Retinal exposure is discussed in a later subsection.

**Table 1.**
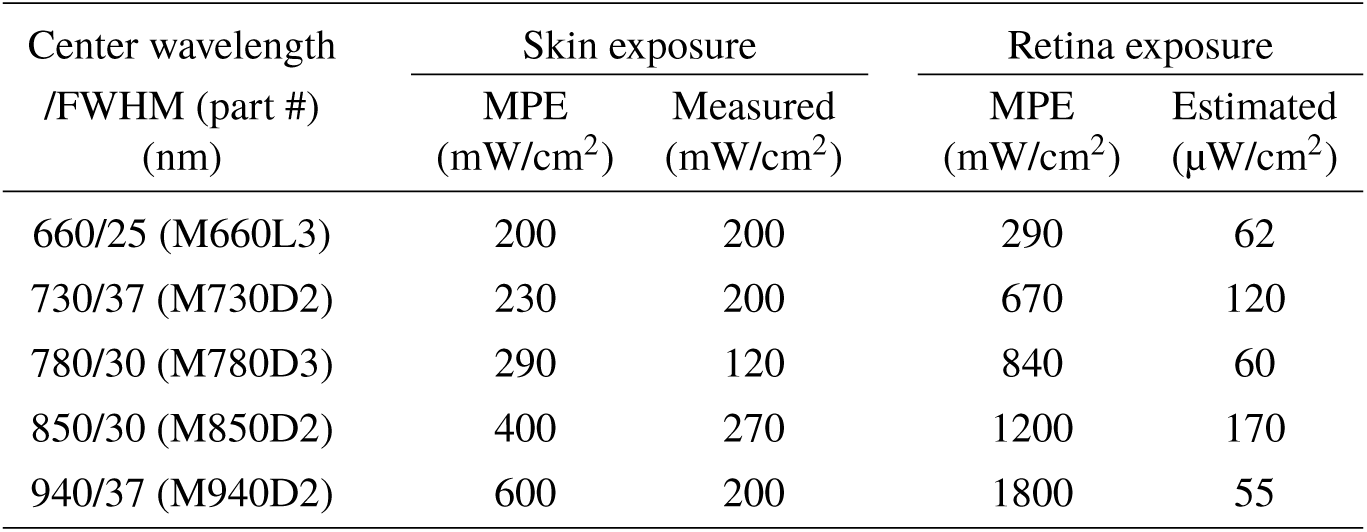
LED choices (with Thorlabs product number), associated ANSI maximum permissible exposures (MPEs) for skin and retina, and measured or estimated irradiance. Retinal MPE is converted from ANSI ocular exposure MPE. Estimation is based on received light power on the camera (see text for details).

| Center wavelength<br>/FWHM (part #)<br>(nm) | Skin exposure |  | Retina exposure |  |
| --- | --- | --- | --- | --- |
|  | MPE<br>(mW/cm<sup>2</sup>) | Measured<br>(mW/cm<sup>2</sup>) | MPE<br>(mW/cm<sup>2</sup>) | Estimated<br>(μW/cm<sup>2</sup>) |
| 660/25 (M660L3) | 200 | 200 | 290 | 62 |
| 730/37 (M730D2) | 230 | 200 | 670 | 120 |
| 780/30 (M780D3) | 290 | 120 | 840 | 60 |
| 850/30 (M850D2) | 400 | 270 | 1200 | 170 |
| 940/37 (M940D2) | 600 | 200 | 1800 | 55 |

#### 2.1.2 Fundus camera and CCD sensor detection subsystem

A modified Topcon TRC-NW5S non-mydriatic fundus camera was used to image the transilluminated fundus. In the present study, the 45°angle of view setting was used exclusively. This angle of view corresponds to a 13-mm arc along the retinal surface. The fundus camera’s built-in illumination system was disabled and a dichroic mirror that normally splits the observation (NIR) and photography (visible flash) beam paths was removed so that all wavelengths collected by the fundus camera were directed to the camera port. The original color camera was replaced with a newer monochrome CCD camera (PCO Pixelfly USB: 1392×1040 pixels, 68 dB dynamic range, 16 ke^-^ well capacity). The camera was operated in “IR boost” mode, which enhances the quantum efficiency (QE, ranging from 47% at 660 nm to 6% at 940 nm).

To make use of the new sensor’s larger format, the total magnification was adjusted by axially displacing the final relay lens and moving the camera to the new focal plane. Final magnification was about 0.5x and camera pixels corresponded to 13 μm on the retina. With additional magnification, pixel size could be further reduced, however, ultimately the numerical aperture (NA), and thus resolution, is limited by the internal stop of the fundus camera itself. For the non-mydriatic fundus camera used in this study, the NA is about 0.038, which in the ideal case yields a diffraction-limited spot size on the retina of about 11 μm for 850 nm light.

To compensate for an intensity gradient from the temporal to nasal side, a gradient neutral density filter (Thorlabs, NDL-25C-2) was inserted near an intermediate image plane and defocused slightly. With higher dynamic range cameras (e.g. sCMOS), this filter is unnecessary. The light transmission of the system was measured using a HeNe laser, co-aligned with the system’s optical axis. Roughly 60% of the beam was transmitted.

#### 2.1.3 Custom GUI and LED toggle

A user interface was written in MATLAB to coordinate and synchronize the illumination and detection subsystems, by way of an Arduino, which was programmed to act as a reconfigurable digital toggle triggered by the camera’s exposure-out signal. Thus, the different LED channels were turned on sequentially in synchrony with the camera exposure.

#### 2.1.4 Transcranial retinal exposure and spectral transmission

The ANSI Z136.1 standard [30] assumes hazardous beams will enter the eye through the pupil. Therefore, the standard specifies ocular MPE in terms of corneal irradiance, which is not directly applicable to transcranial transmission imaging where light primarily exposes the retina from behind. Assuming an “extended source”, the corneal MPEs were converted to equivalent retinal MPEs as described previously [31], and listed in Table 1.

While a direct measurement of the actual in vivo retinal exposure is impossible, with knowledge of several parameters, we may estimate a quantitative relationship between recorded pixel value and retinal exposure. These parameters are camera pixel area (41.6 μm^2^), QE, exposure time, magnification, system transmittance, ocular media transmittance [32], entrance pupil diameter (~1 mm, set by fundus camera), and eye focal length (17 mm). Additionally, we assume the light at the retina has a Lambertian angular distribution. To account for attenuation due to the retinal pigmented epithelium (RPE), we assume the light exiting the retina has also traversed a 20 μm-thick absorbing layer with a wavelength-dependent absorption coefficient [33]. The estimated exposures are all orders of magnitude below the MPE values, which should allay concerns about the absolute accuracy of this estimate.

Since LEDs have moderate spectral bandwidth (see Table 1), transmission through skin, bone, and brain is expected to only moderately impact the shape of the spectrum incident on the back of the fundus. This was verified separately by temporarily replacing the camera in our system with a fiber-coupled spectrometer (Thorlabs CCS175 with 1 mm-diameter fiber patch cable). The empirical transmission results were combined with the QE and the resultant transmission-responsivity curve was used to correct the weighted average absorption coefficient for each chromophore (HbO_2_ and Hb) at each LED channel. The weighted average absorption coefficients in the intermediate wavelength range (730, 780, and 850 nm LEDs) were found to be less sensitive to the correction (<2% change). However, at 660 nm the weighted average absorption coefficients of HbO_2_ and Hb were each reduced by 8%. This is primarily due to strong absorption on the shorter wavelength end of the LED spectrum. The light spectrum incident on the retina is red-shifted, and hence the absorption coefficients are reduced since the spectra are both decreasing around 660 nm. Additionally the weighted average absorption coefficient of deoxyhemoglobin was increased by 26% at 940 nm. In this case the dramatic change is mostly due to the dwindling QE around 940 nm, which effectively blue-shifts the detectable spectrum.

### 2.2. Subjects and imaging sessions

Four asymptomatic eyes from four normal subjects (23–59 years of age, mean age 46, one female and three males) were imaged. The subjects presented a wide range of choroidal pigmentation, with corresponding (usually correlated) iris colors ranging from blue to brown. For each subject, informed consent was obtained prior to imaging. The research was approved by the Boston University Institutional Review Board.

For each imaging session, the subject fixated on an external target such that the optic disc was centered in the image. Although the system has the capability to temporally interleave LEDs for quasi-simultaneous multispectral imaging, the camera’s maximum framerate limited this capability. For instance, enabling three color channels reduced the multispectral framerate to 4 Hz. Because light from the 660 and 730 nm could be weakly perceived, while the others LEDs generally could not, temporal multiplexing of the LEDs effectively caused 4 Hz blinking across the whole field of vision, which was irritating to the subject. Instead, a long series of frames from the same LED were acquired. A 256-frame series was completed in about 26 sec. For one subject (59-year-old white male, moderate pigmentation), a complete multispectral dataset was acquired by repeating this process for all five LEDs. This dataset was chosen for more detailed analysis and displayed throughout this article.

To ensure that the ANSI-equivalent retinal MPE was not exceeded, the MPE-equivalent maximum allowable pixel value (for a given exposure time) was calculated a priori. The LED current was carefully increased and pixel values were continually monitored to confirm permissible exposure. None of the light conditions used in this study came close to exceeding equivalent retinal MPEs.

### 2.3. Image processing

Although the use of NIR light permits deep tissue penetration, its major disadvantage is significantly reduced intrinsic absorption contrast, particularly for hemoglobin, which drops precipitously past 600 nm [34]. For example, consider a 30 μm-diameter arteriolar vessel (about the size of the small vessel labeled with arrowheads in Fig. 2B & C). The absorption coefficient of highly oxygenated arterial blood at 850 nm is approximately 5.5 cm^-1^. Light passing through the diameter of this vessel is only attenuated 1.6%. With the exposure conditions used to obtain Fig. 2A, an average of 3800 photoelectrons (e^-^) per pixel are detected. The expected signal due to the small vessel is thus 61 e^-^. In comparison, the rms noise in a shot noise-limited system is 62 e^-^, meaning that the signal to noise ratio (SNR) is no better than unity. Therefore, the vessel cannot be reliably detected with the exposure conditions used for Fig. 2A.

**Fig. 2.**
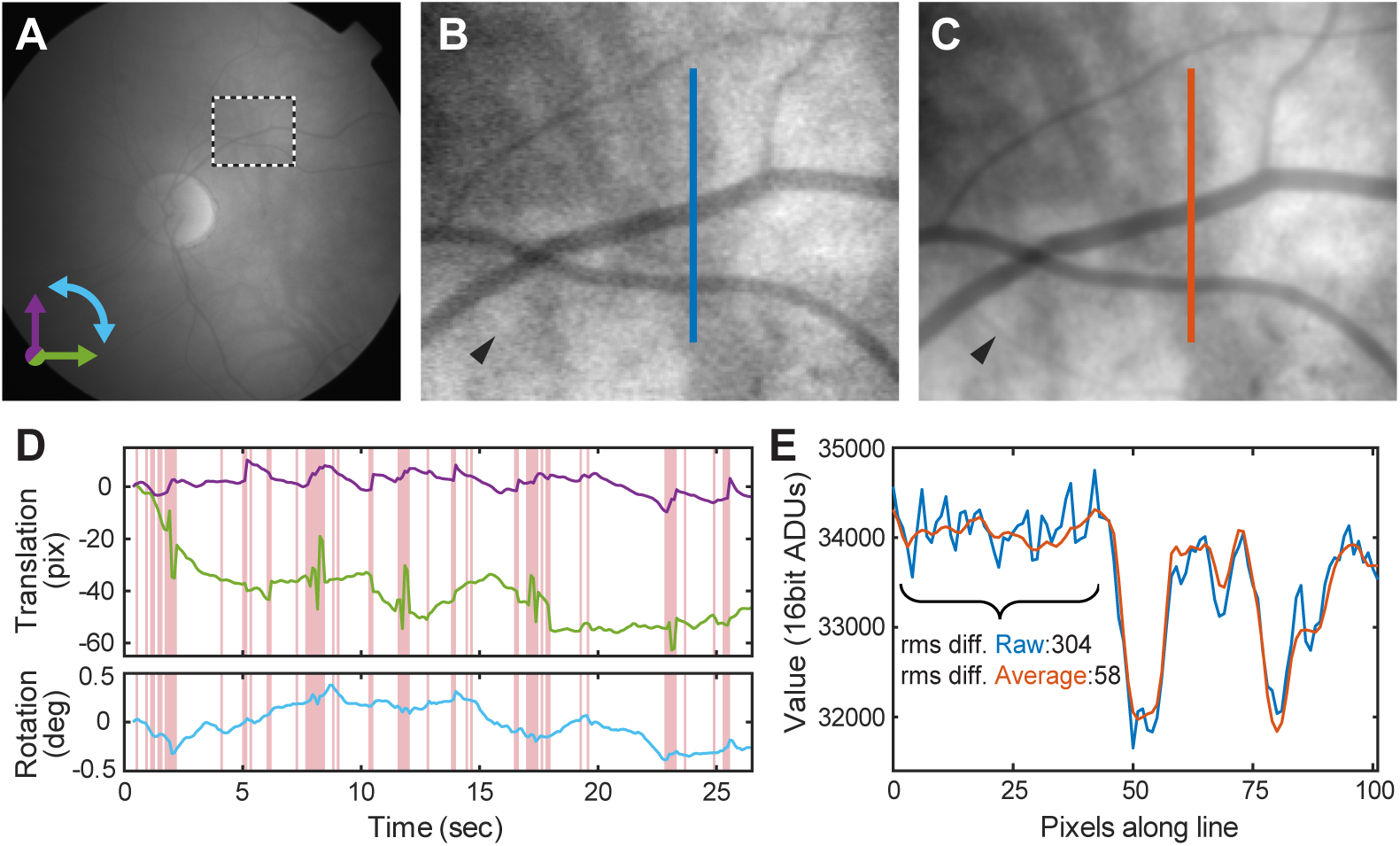
Registration and averaging. A: Single frame from 850 nm LED image series (10 Hz acquisition). B: Magnified view of highlighted region in A. Black/white levels are scaled to the region’s min/max values. C: 194-frame average improves SNR, revealing previously undetectable small features (arrowheads in B & C). D: Detected translation (green: X, magenta: Y) and rotation (cyan) for 256-frame series. Motion-corrupted frames (red-background) are excluded from averaging. E: Profiles for lines drawn in B & C, demonstrating SNR improvement. (rms diff.: Root-mean-square pixel to pixel difference)

#### 2.3.1. Image registration and averaging

To improve SNR (i.e. increase detected photoelectrons), we record a series of shot-noise-limited frames, register each frame post-hoc, and average the registered frame series in time. Custom software was written in MATLAB, based on refs. [35,36], to register sequential frames to subpixel precision and account for slight rotations from torsional eye movements. A plot of the detected translational (green: X, magenta: Y) and rotational (cyan) movements versus time for a typical image series is shown in Fig. 2D. With the exposure duration used in the present study (~100 ms), eye motion inevitably corrupts a number of frames. The registration software detects and automatically excludes motion-corrupted frames from the final high-SNR average (shown as red background in Fig. 2D). The number of frames averaged ranged from 125 to 223 (out of 256), depending on the level of eye movement. The camera records 14-bit images, however we expand this range to 16-bits for data storage.

A comparison between a single frame and the high-SNR average frame is shown in Fig. 2B & C with attendant line profiles plotted in Fig. 2E. No apparent loss of resolution is observed and variation is now limited by the underlying variation of the fundus itself.

#### 2.3.2. Relative absorbance calculation

Absorbance is calculated from transmitted and incident light power measurements: *A* = −log(*I*_trans_/*I*_inc_). As is apparent in Fig. 2A, the illumination for transmission fundus imaging was not spatially uniform. To accurately compute absolute absorbance, the incident light power as a function of location must be known. Since direct measurement of this is infeasible, we instead calculate a relative absorbance as follows

First, the high-SNR average image (Fig. 3A) is corrected for slowly varying global illumination changes by fitting the image to a polynomial surface, avoiding the bright optic disc area. The average image is divided by the fit to produce a flattened image (Fig. 3B). Next the flattened image is local maximum filtered, that is, each pixel is replaced with the maximum value in a circular local area around the pixel. The filtered image is smoothed with a Gaussian kernel equal to half the diameter of the local maximum filter, producing Fig. 3C. This smoothed local maximum image is used as a proxy for the incident light power to compute absorbance. In other words, this method calculates absorbance *relative the minimum absorbance* in a local neighborhood around each pixel. Technically, this should be referred to as relative attenuation, a more general term that additionally includes loss due to scattering outside the acceptance angle of the optical system. Such loss may happen, for instance, at blood vessels where erythrocytes are known to cause scattering [37]. However, in subsequent analysis the effect of spatially varying scattering loss is neglected. The assumption is that scattering from the choroid to the cornea is reduced and/or very strongly forward-directed when using NIR light.

**Fig. 3.**
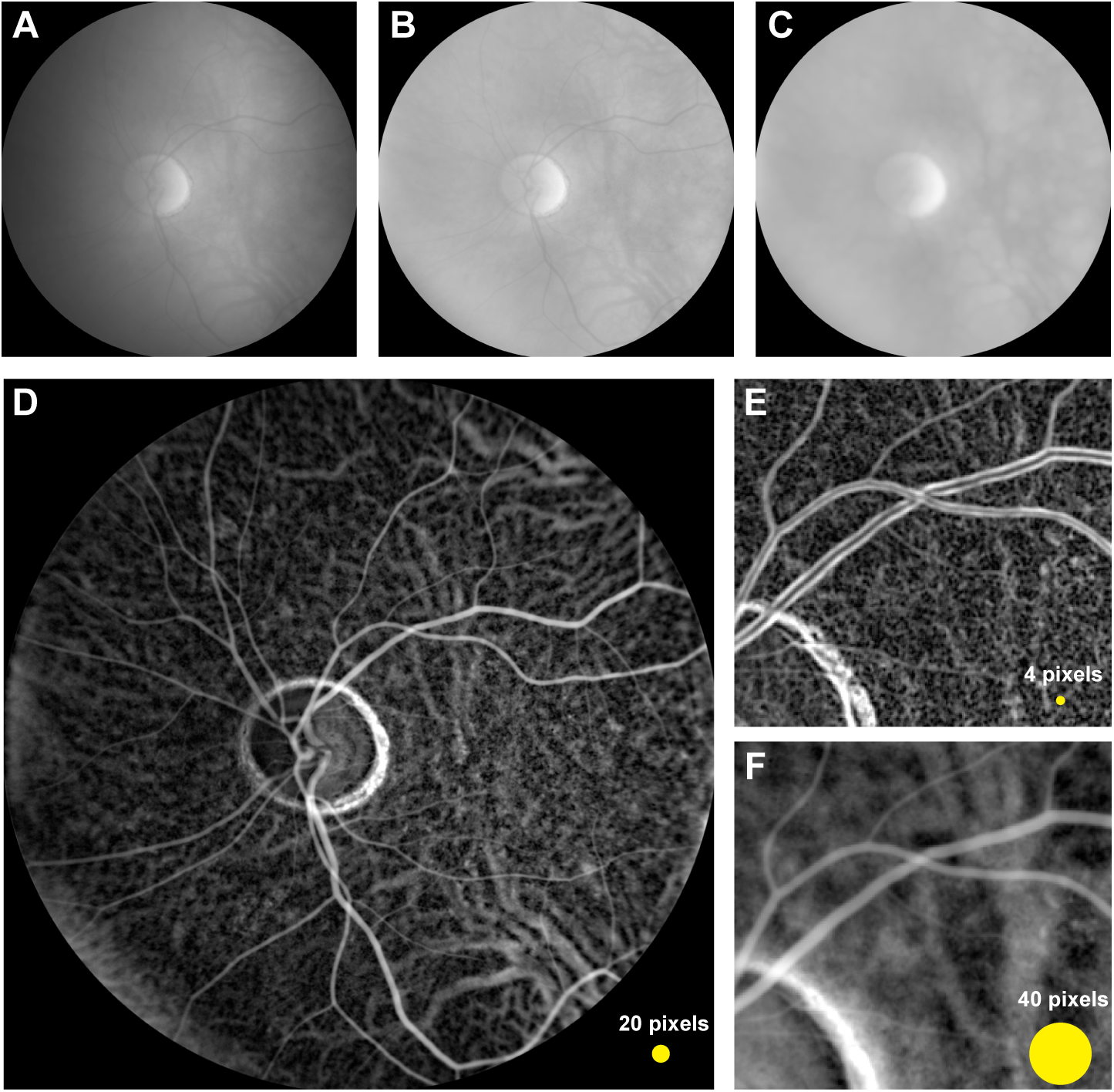
Left eye from a 59-year-old healthy subject. A-D: Relative absorbance calculation with 850 nm transillumination (refer to text for details). Yellow circle indicates size of the local maximum filter kernel relative the image. E & F: Effect of changing local maximum filter kernel radius (zoomed region of D). A smaller radius emphasizes sharp changes in absorbance, revealing mottled pigmentation density fluctuations. Larger kernels emphasize larger features such as vessels in the choroid.

## 3 Results

As predicted, the transillumination images in Figs. 2–3 are free from spurious back-reflections, either from the inner limiting membrane or nerve fiber layer (NFL), which, in a conventional reflection-mode imaging, collectively appear as a glare that intensifies towards the optic disc. The images presented here (Fig. 2B & C, especially) are also free from central vessel reflex. Interestingly, the optic disc appears bright compared with the fundus background, possibly due to anisotropic forward scattering parallel the axonal fibers, and also likely due to the absence of underlying choroid and RPE pigment.

The contrast of the relative absorbance image (Fig. 3D) resembles that obtained with fluorescein angiography, since at 850 nm HbO_2_ and Hb have similar absorption. A bright ring around the border of the optic disc is generated when the local maximum filter’s spatial kernel intersects with the (relatively) bright disc. Thus the filter overestimates incident light power near the optic disc. A more sophisticated approach might segment the disc from its surroundings and compute local maxima for each domain separately. Regardless, we caution against using the relative absorbance representation when inspecting interfaces, such at the edge of the optic disc or, in cases of AMD or diabetic retinopathy, near retinal scars, edemas, or focal deposits. Instead we recommend using the transmitted image (Fig. 3A or B), which should be free from this processing artifact.

In the relative absorbance images, visibility of certain features appears to depend on the diameter of the maximum filter kernel used. Namely, a small kernel highlights rapidly varying, tenuous features, tantamount to high-pass filtration because the local-maximum-filtered image over such a small kernel is nearly a low-pass version of the image. Outside the optic disc, the distinctive mottled appearance of RPE pigment density variations is apparent [10]. On the other hand, larger kernels emphasize large-scale features, such as large choroidal vessels, which are clearly visible in the background of Fig. 3F.

Notably, there is no evidence of the macula, which under visible light fundus photography ordinarily appears as a dark spot 2–3 optic disc diameters temporal the optic disc. The absorbance of macular xanthophylls is well-known to be low for the NIR wavelengths used in this study [38].

### 3.1 Multispectral imaging and chromophore unmixing

Data obtained using the other LED channels are processed in a similar way and shown in Fig. 4. As the wavelength of transillumination is scanned, two changes are obvious. First, for shorter wavelengths (660 and 730 nm), retinal arteries nearly disappear. This finding is well-known and explained by noting that under normoxia, arteries are highly oxygenated, and the absorbance of oxygenated hemoglobin is minimal around 690 nm (see Fig. 5A for complete HbO_2_ and Hb absorption spectra). Retinal veins, however, contain a mixture of HbO_2_ and Hb, which results in broadband absorbance across the NIR spectrum and explains why retinal veins are visible in each spectral image.

**Fig. 4.**
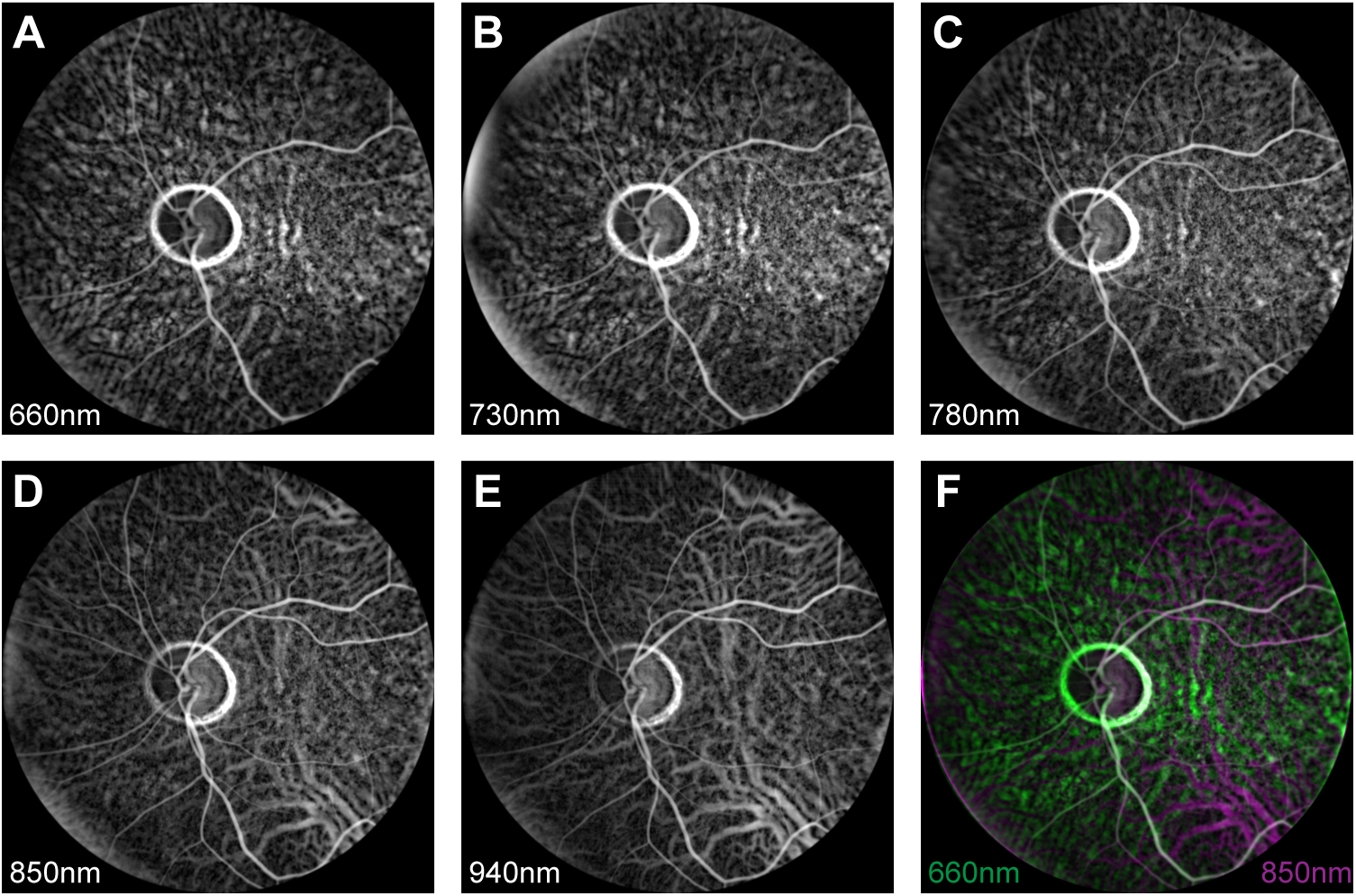
A-E: Multispectral relative absorbance images. Brightness has been autoscaled for clarity. F: 660 and 850 nm images overlaid in false-color for comparison. Vortex veins are readily observed in the top right and bottom right fields of 850 and 940 nm images.

The second notable change involves a contrast inversion of the large choroidal vessels. For shorter wavelengths, choroidal vessels appear dark, meaning that they are weaker absorbers than their surroundings (i.e. the extravascular stroma). For longer wavelengths (850 and 940 nm), choroidal vessels appear bright, meaning that they absorb more than the extravascular stroma. The extravascular stroma contains melanocytes which, for short wavelengths (660 and 730 nm), are highly absorbing. With increasing wavelength, the absorption of melanin monotonically decreases (Fig. 5A, dotted line), while oxyhemoglobin absorption increases. Since choroidal vasculature is known to be highly oxygenated, beyond 780 nm, the choroidal vessels exhibit more absorbance than the extravascular melanin-containing stroma, explaining the contrast inversion. These observations are consistent with previous studies [10, 39] that assessed visibility of subretinal structures with NIR light.

**Fig. 5.**
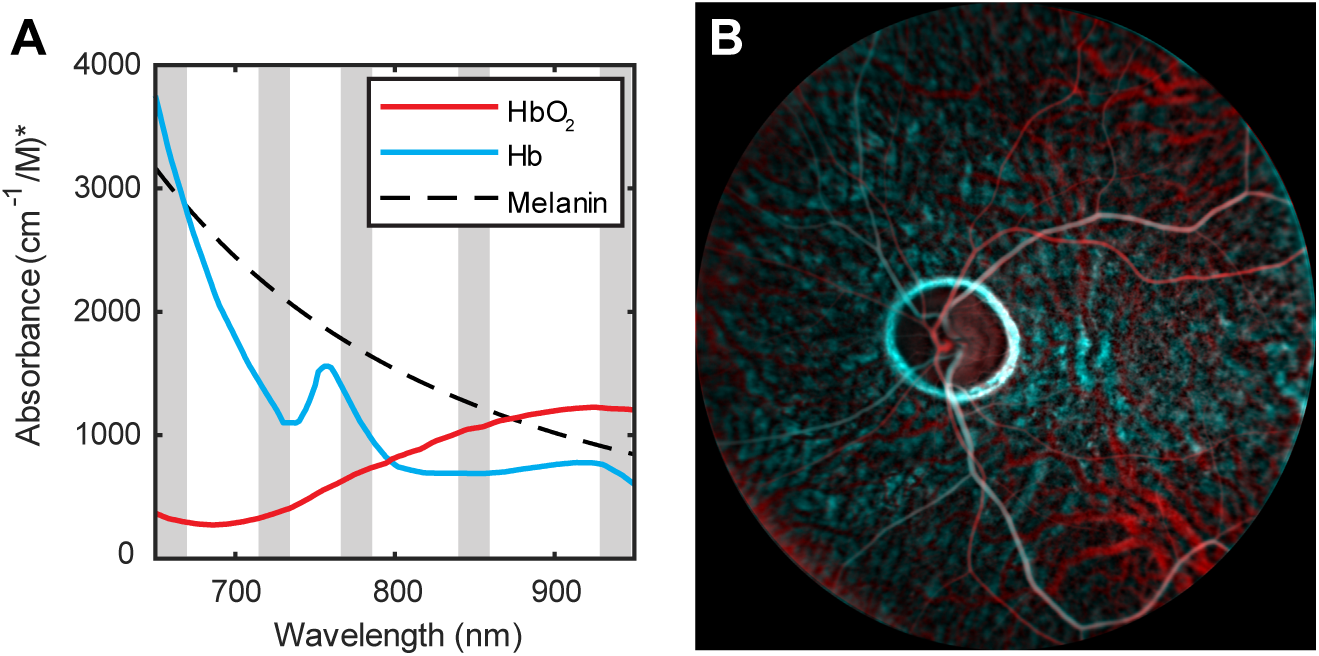
A: The absorption spectra for the three dominant NIR chromophores of the eye. Gray background indicates available LED channel. B: Oxy- and deoxyhemoglobin (HbO2 and Hb) distributions based on absorption spectra in A and non-negative least squares spectral unmixing, using 660-850nm relative absorbance images. Hemoglobin spectra obtained from Ref. [34] and average melanin slope from [33]. (*Melanin spectrum scaled to permit comparison with hemoglobin. Units do not apply.)

To register each spectral image, we manually selected at least 10 retinal vein bifurcation points, estimated the movement, and applied a similarity transform. Two registered images, which exhibit both contrast changes mentioned above, are shown together in false color in Fig. 4F.

With spatially aligned spectral images, one can also “unmix” the contributions from individual chromophores to the total observed absorbance via the Beer-Lambert law. Thus, a system of equations relating chromophore concentration to absorbance at each spectral channel may be written. Concentrations are averages over the volume approximately defined by the depth of focus and pixel area. The coefficients are equal to the weighted-average absorption coefficients for each chromophore over each LED spectrum (corrected with the transmission-responsivity curve explained in Section 2.4.1). The system of equations is solved separately at each pixel using either a pseudoinverse or a non-negative least squares optimization routine (standard functions in MATLAB), the latter applying a positivity constraint to the estimated chromophore concentrations. The non-negative least squares solution was preferred for displaying chromophore maps, but produced less smoothly varying background estimates, particularly when the estimated chromophore concentration was around zero. The pseudoinverse solution yielded a smooth background which was preferable for the image inpainting problem described later.

Non-negative least squares-based unmixing of HbO_2_ and Hb is shown in Fig. 5B. The 940 nm channel was not included due to uncertainty in the large 26% weighted-average absorption coefficient correction for deoxyhemoglobin. The deoxyhemoglobin channel (cyan) in Fig. 5 likely contains contribution from melanin in the pigmented extravascular choroidal stroma. However, incorporating melanin as a third component in unmixing yielded poor results (not shown), likely because at the discrete wavelengths used, the orthogonality between Hb and melanin is small.

### 3.2 Retinal oximetry

With estimates for HbO_2_ and Hb concentration, oxygen saturation, SO_2_ = [HbO_2_]/([Hb] + [HbO_2_]) may be readily calculated. For a quantitative comparison with results from previous retinal oximetry studies, chromophore concentrations in retinal vessels must be isolated from the influence of the choroidal background, which can be quite significant for NIR transillumination. Relative to the retinal vessels, the background is generally slowly varying, (i.e. only the largest choroidal vessels are visible). Thus the background may be numerically estimated using an image restoration technique known as inpainting. For this study, a simple inpainting algorithm that is included in MATLAB was used. Essentially the algorithm interpolates unknown pixel values based on the solution to Laplace’s equation subject to discrete 2D boundary conditions given by the value of the surrounding pixels. The process is graphically illustrated in Fig. 6A. First, clearly delineated retinal vessels are manually segmented from HbO_2_ and Hb maps. Pixels within segmented vessels are set to zero. The background is estimated with inward interpolation (inpainting). Finally, the background-estimated image is subtracted from the original chromophore image, which removes substantial variation and offset introduced by the choroidal background.

**Fig. 6.**
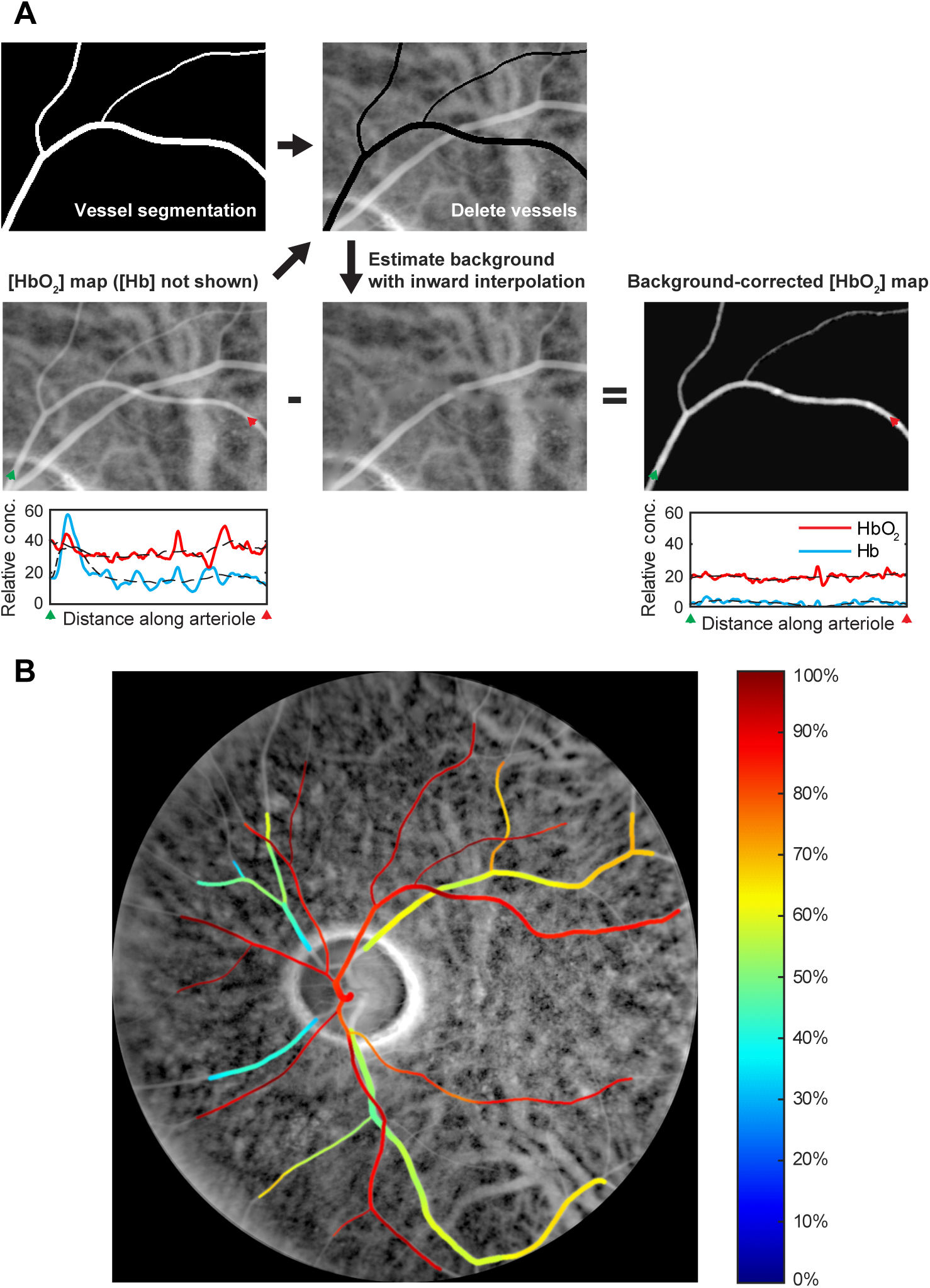
Retinal oximetry in fundus of the left eye of a normal volunteer. Input data is spectrally unmixed HbO_2_ and Hb relative absorbance images using the psuedo-inverse solution. A: Processing workflow leading to background-corrected vessel chromophore maps (refer to text for explanation). B: Pseudocolor overlay of oxygen percent saturation (SO_2_) in major retinal vessels.

For retinal oximetry, a larger local maximum filter kernel (32 pixels) was used for the relative absorbance calculation and the pseudoinverse was used to provide the spectral solution. These tended to produce a more smoothly varying background for which the background removal strategy outlined above performed well. Several large branching retinal arteries and veins were segmented and background-corrected. Veins over the optic disc exhibited mild spontaneous venous pulsation causing blurring in the average image. These veins segments were therefore excluded from oximetric analysis. The corrected vessel maps were smoothed using a low pass filter (Gaussian, 32-pixel FWHM), to remove disruptions near vessel intersections. The effect of this low pass-filter is shown as dotted lines in the line profiles of Fig. 6A. Estimated SO_2_ is overlayed in pseudocolor onto a map of total hemoglobin concentration and shown in Fig. 6B.

Additionally, the mean and standard deviation of SO_2_ were computed for different anatomical regions around the optic disc and summarized in Table 2. The arteriovenous (A-V) SO_2_ difference is also given for each region, as a relative indicator of local oxygen extraction and metabolic demand. The combined average SO_2_ was 89% for arteries and 57% for veins (32% A-V difference).

**Table 2.** Quantified retinal oximetry results (% oxygen saturation, SO_2_) in regional quadrants and across the whole fundus shown in Fig. 6 (temporal: right, superior: top).

| Anatomical region | SO<sub>2</sub> mean ± S.D. (# pixels) | | A-V difference |
| --- | --- | --- | --- |
|  | Arterial | Venous |  |
| Superior temporal | 88 ± 5 (6849) | 64 ± 4 (5539) | 24 |
| Superior nasal | 92 ± 6 (3229) | 48 ± 7 (2543) | 44 |
| Inferior nasal | 93 ± 4 (1097) | 39 ± 3 (1404) | 54 |
| Inferior temporal | 87 ± 6 (4877) | 58 ± 5 (7139) | 29 |
| Combined | 89 ± 6 (16,761) | 57 ± 9 (16,625) | 32 |

## 4. Discussion

Compared with prior work, the arterial and venous retinal SO_2_ values presented here appear reasonable. Previously, for arterial SO_2_, Delori reported 98% using a scanning three-visible wavelength approach [40]. De Kock et al. reported 97% with a time-resolved whole fundus pulse oximeter [41]. Hyperspectral approaches yielded 92% [18] and 104% [42], and recently visible light OCT reported an average across subjects of 92% [43]. For venous SO_2_, Hickam et al. reported 59% at the optic disc using a two-color photographic method [21]. Delori reported 45% [40]. Beach and colleagues used a digital two-color imaging method to calculate 55% after correcting for pigmentation and vessel diameter effects [22]. Hyperspectral techniques yielded 35% [42] and 58% [18], while an average of 77% was found using visible light OCT [43].

For this subject, the nasal quadrants exhibited greater A-V differences. Interestingly, this is contrary to the findings of Schweitzer et al. [18], who reported a statistically significant increase in A-V difference in the inferior temporal quadrant compared with the inferior nasal quadrant. Since retinal oxygen consumption is greater in darkness [44], one possible explanation might be that the nasal regions of the transilluminated retina, which are farther away from the light source, are exposed to less light, thus their oxygen consumption is increased relative to the temporal side. This effect would only be significant for the 660 nm LED channel, however this channel plays an outsized role in spectral unmixing due to the large difference in absorption between HbO_2_ and Hb. It is also possible that the gradient neutral density filter may not be perfectly neutral, leading to a system-induced bias on one side. More transillumination imaging data (preferably without the filter) is needed to confirm this relation.

It should be stressed that the oximetric results presented here were obtained without external calibration and instead used only the published extinction spectra for HbO_2_ and Hb. An external calibration is time-consuming, but useful in accounting for attenuation or spurious back-reflections due to tissue scattering [22]. In this study, it was assumed that NIR light was weakly and predominantly forward-scattered, which is a generally valid for biological tissue. Thus, local attenuation due to spatially varying scattering (e.g. at blood vessels) was ignored and relative concentrations of oxy- and deoxyhemoglobin were calculated according to the Beer-Lambert law. This assumption is probably not absolutely true since it is known that whole blood deviates from Beer-Lambert due to multiple scattering [45]. The attenuation due to scattering may also falsely contribute to the unmixed deoxyhemoglobin, since its spectral dependence is likely to be similar to the absorption of deoxyhemoglobin, which could explain why the average arterial SO_2_ determined here is somewhat lower than previous results and also why incorporating melanin into the spectral unmixing led to unsatisfactory results. Monte Carlo simulations of fundus transillumination may provide useful insights and help resolve the question of scattering attenuation in this new geometry.

Another obvious question concerns the validity of spectroscopy obtained over the course of a few minutes. Because our camera’s low framerate prevented temporally interleaved multispectral imaging, the different LED channels were recording sequentially over a total period of 2 minutes. It is possible that vessel oxygenation can vary over this time period, however Hickham and colleagues have found little SO_2_ variation over this time period [21], so such dynamics may indeed be safely neglected.

Compared with reflection-based fundus photography, transillumination fundus imaging offers several unique advantages. First, imaging penetration is deep, meaning that discrete choroidal vessels are plainly visible. This is due to a combination of the NIR wavelengths used, which are only weakly absorbed and scattered in the inner retina, and the absence of spurious back-reflections, which can easily overwhelm weak signal emanating from deeper structures. Elsner and colleagues [39] have exploited the same factors to image sub-retinal structures, but rather than a transmission geometry, they used confocal detection and an annular aperture to reject inner retinal reflection. There are also now OCT-based approaches for deep choroidal imaging [46], which intrinsically separate anterior surface scatter. Compared to these techniques, transillumination instrumentation is considerably less complicated, and thus may be more appropriate for high throughput disease screening. Also because the technique is single-pass, transillumination may offer some advantage in imaging retinas of patients with cataracts or other opacities.

Second, transillumination ensures that all light detected has traversed the tissue or vessel of interest. This has important implications for retinal oximetry, where vessel surface reflections are a common confounding factor [13]. Researchers have attempted to compensate for these reflections either by fitting the wings of the vessel profile to a Gaussian model [47] or by choosing to analyze only the darkest point along the profile [23]. Both of these techniques reduce the sensitivity because they essentially discard possible data points. In contrast, transillumination is free from surface reflections and may utilize the entire vessel profile for oximetric calculations.

The proof-of-concept system detailed here is far from optimized. The SNR, or more precisely the absorption contrast to noise ratio, is primarily limited by shot noise, which is fundamentally linked to the number of photoelectrons generated in the camera sensor. The adoption of newer CMOS sensors, which feature higher NIR sensitivity (up to 2.5x greater at 850 nm) and faster readout times, will increase the number of possible detected photoelectrons. As evident in Table 1, there is also the possibility of using more power, by adding several of the same LED and/or spreading the incident light over a larger area of skin. Illumination from other locations, such as the palate or nose, may also offer improved or complementary transillumination efficiency. For longer wavelengths, significantly more power may be used before reaching the ANSI Z136.1 skin MPE. This is fortunate since the quantum efficiency of silicon-based cameras decreases sharply in this spectral range. Since our aim is to demonstrate fundus transillumination on a clinically compatible platform, the current system was designed around a commercial non-mydriatic fundus camera. However, the fundus camera’s internal stop limits detection to just 1 mm of pupil diameter. If the fundus camera is instead replaced with a system capable of utilizing the entire naturally dilated pupil (up to 8 mm) for detection, light throughput is expected to increase by a factor of 64, although likely at the expense of resolution.

In conclusion, an alternative illumination method for human non-mydriatic fundus imaging has been presented. It is based on the transcranial delivery of NIR light and on multiple scattering to redirect a portion of this light to the posterior eye, resulting in glare-free chorioretinal imaging. The use of NIR light enables transillumination, however at the expense of absorption contrast. We showed that with careful image processing, the contrast to noise ratio may be rendered adequate for oximetry measurements. Compared with conventional reflection-based fundus photography techniques, NIR transillumination simplifies absorption measurements and allows imaging deep into the choroid. Importantly, the technique is compatible with reflection-based techniques and we have shown that it works well with a commercial fundus camera. Combining information from these two illumination approaches may improve spectroscopic analysis of the fundus.

## Funding

Boston University Photonics Center

## Acknowledgments

The authors are indebted to Thomas Bifano and the Boston Micromachine Corporation for providing the fundus camera used in this study. TDW also wishes to thank David Boas and Charles Lin for helpful discussions and encouragement.

## Disclosures

The authors declare that there are no conflicts of interest related to this article.

